# Changes in m^6^A RNA methylation of goat lung following PPRV infection

**DOI:** 10.1101/2022.03.24.485342

**Authors:** Raja Ishaq Nabi Khan, Manas Ranjan Praharaj, Waseem Akram Malla, Neelima Hosamani, Shikha Saxena, Bina Mishra, Kaushal Kishor Rajak, Muthuchelvan Dhanavelu, Ashok Kumar Tiwari, B P Mishra, Basavaraj Sajjanar, Ravi Kumar Gandham

## Abstract

Peste des petits ruminants (PPR) is an acute, highly contagious viral disease of goats and sheep, caused by the Peste des petits ruminants virus (PPRV). Earlier studies suggest the involvement of diverse regulatory mechanisms in PPRV infection. Methylation at N6 of Adenosine called m^6^A is one such RNA modification that influences various physiological and pathological phenomena. As the lung tissue represents the primary target organ of PPRV, the present study explored the m^6^A changes and their functional significance in PPRV disease pathogenesis. m^6^A-seq analysis revealed 1289 m^6^A peaks to be significantly altered in PPRV infected lung in comparison to normal lung, out of which 975 m^6^A peaks were hypomethylated and 314 peaks were hypermethylated. Importantly, hypomethylated genes were enriched in Interleukin-4 and Interleukin-13 signaling and various processes associated with extracellular matrix organization. Further, of the 843 differentially m^6^A containing cellular transcripts, 282 transcripts were also found to be differentially expressed. Functional analysis revealed that these 282 transcripts are significantly enriched in signaling by Interleukins, extracellular matrix organization, cytokine signaling in the immune system, signaling by receptor tyrosine kinases, and Toll-like Receptor Cascades. We also found m^6^A reader HNRNPC and the core component of methyltransferase complex METTL14 to be highly upregulated than the m^6^A readers – HNRNPA2B1 and YTHDF1 at the transcriptome level. These findings suggest that alteration in m^6^A landscape following PPRV is implicated in diverse processes including the Interleukin signaling.

## Introduction

PPR is an acute, highly contagious viral disease of goats and sheep, caused by the Peste des petits ruminants virus (PPRV), a morbillivirus in the family Paramyxoviridae [1]. PPR is characterized by mucous membrane congestion, pyrexia, pneumonia, nasal, ocular discharge, diarrhea, necrotic stomatitis, and leukopenia [2]. The disease is highly prevalent across sub-Saharan Africa, the Arabian Peninsula, and the Indian subcontinent [3]. Morbidity and mortality rates can reach 90-100% leading to severe economic losses in the endemic regions [4]. Consequently, PPR has been classified as a World Organization for Animal Health (OIE) listed disease. The causative agent is a non-segmented RNA virus with a genome size of 15,948 nucleotides that encodes six structural proteins [nucleocapsid (N), phosphoprotein (P), matrix protein (M), fusion protein (F), haemagglutinin protein (H), and large protein (L)] in 3’ to 5’ direction and two non-structural proteins V and C resulting from the RNA editing of phosphoprotein (P) [1].

Various studies have been carried out to understand PPRV-host interactions, since the advent of next-generation sequencing, We previously studied the effect of the PPRV vaccine strain (Sungri/96) on goat PBMCs at the transcriptome level and reported a significant alteration in the expression of the genes involved in the regulation of innate immunity and apoptosis [5]. The dysregulation of innate immunity during PPRV infection of goat PBMCs has also been reported by a different group [6]. In a virulent PPRV-host interactions study, lymphocytes have been observed to contribute more to counterattacking the PPRV infection whereas the contribution of monocytes was found to be limited [7]. The expression levels of viral sensors and interferon-stimulated genes were found elevated only in lymphocytes following the viral infection [7]. Besides, the role of miRNAs in regulating PPRV infection has been reported[8, 9]suggestive of the involvement of diverse regulatory mechanisms in PPRV infection.

Epitranscriptomics (=RNA epigenetics) involves functionally relevant biochemical modifications of RNA. More than 150 types of RNA modifications are present in different RNA types and among them, m^6^A is most abundant in mRNAs and functionally well characterized [10, 11]. The enzymes that install the m^6^A modification (=m^6^A writers), that remove it (=m^6^A erasers), and certain RNA binding proteins (=m^6^A readers) that affect the fate of the m^6^A-containing RNA by recruiting various proteins, have been found. m^6^A influences RNA metabolism, which in turn influences various physiological and pathological conditions [12]. To identify the m^6^A sites at the transcriptome level, antibody-based screening methods such as methylated RNA immunoprecipitation sequencing (meRIP-seq/m^6^A-seq) are being widely used [13, 14]. RNA epigenetic changes are novel regulatory mechanisms that are strongly implicated in virus-host interactions. Zika Virus has been found to induce changes in m^6^A modification of host RNA that affect translation, splicing, and mRNA stability [15]. Similarly, HIV-1 Infection of T-cells has been found to influence the host mRNA methylomes [16, 17]. KSHV(Kaposi’s sarcoma-associated herpesvirus) latent infection regulates tumorigenic and epithelial-mesenchymal transition pathways by causing hypomethylation within 5′ untranslated region (UTR) and hypermethylation with 3′ UTRof host RNA [18]. There have been no studies on the role of RNA modifications in the pathogenesis of PPRV in small ruminants.

The histological examination of the lungs of PPRV infected goats reveal proliferative changes like bronchitis, bronchiolitis, the proliferation of type II pneumocytes, and the presence of large multinucleated syncytial giant cells [2, 19, 20]. As the lung tissue represents the primary target organ of PPRV, the present study explored the epitranscriptomic changes and their functional significance in PPRV disease pathogenesis.

## Materials and methods

### Ethical Statement

The animals (n=2) used for the study were taken from the Indian Veterinary Research Institution (IVRI). The permission to conduct the study was granted by Institutional Animal Ethics Committee (IVRI-IAEC) under the Committee for Control and Supervision of Experiments on Animals (CPCSEA), India, vide letter number F.26-2/2019/JD(R) dated 22^nd^ April 2020. All relevant guidelines were followed for the study.

### Experimental animals and Sample collection

An overview of the experiment is shown in **Figure 1**. The experimental infection is part of the vaccine potency testing experiment. Goats (1 year of age) for the experiment were initially tested to be negative for the presence of PPRV antibodies by competitive ELISA [21]and serum neutralization test [22]. A highly virulent PPRV (Izatnagar/94 - lineage IV; [23]) isolate maintained at PPR Laboratory, Division of Virology, Indian Veterinary Research Institute, Mukteshwar was used as a challenge virus. A splenic suspension (10%) of the virulent virus was inoculated subcutaneously (4 ml suspension, 2 ml each at two different sites). The infected animals were sacrificed after they manifested the characteristic clinical symptoms of PPRV infection and lung tissue samples were collected [24]. For this comparative study, lung samples of apparently healthy goats of the same age group were obtained from the nearby slaughterhouse as well. All the samples were stored in RNAlater RNA stabilizing solution(Qiagen, #76106) at 4°C and then stored at – 80°C before proceeding further.

**1.**
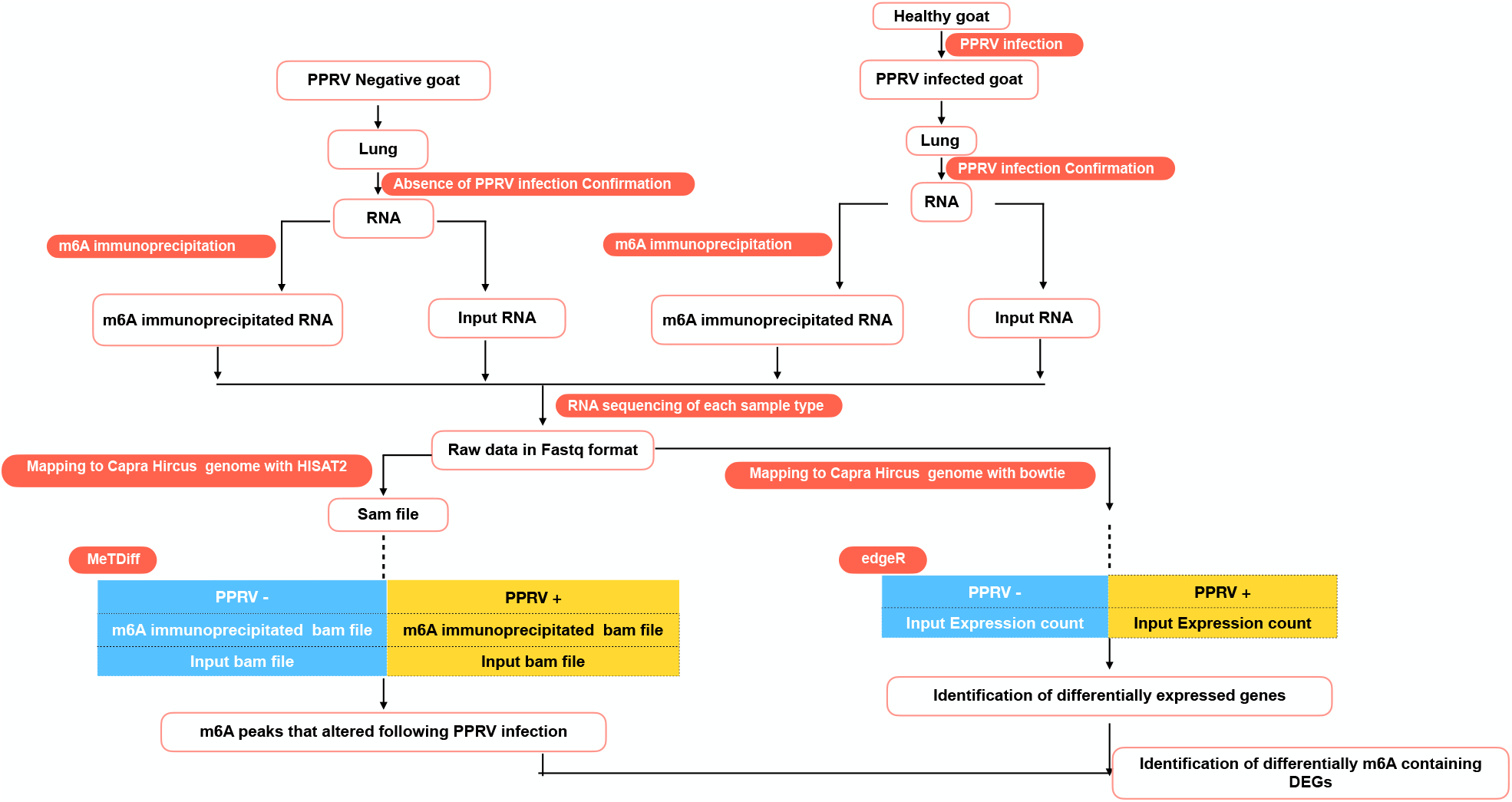
Workflow for m^6^A sequencing analysis and RNA sequencing analysis.

### Sandwich ELISA (s-ELISA)

The samples were initially triturated in PBS followed by two cycles of freezing and thawing. s-ELISA of the supernatant collected was performed as per the instructions provided with the kit [25]. Briefly, 100 μl capture antibody diluted in the ratio of 1:4000 was taken and then dispensed in all the wells of an ELISA plate. The ELISA plate was kept in the incubator on an orbital shaker at 37°C for incubation up to 1 h. At the end of incubation, the unbound antibodies were removed using the wash buffer composed of PBS and Tween-20. This step was repeated thrice. Thereafter, 50 μl of blocking buffer (0.01M PBS containing 0.1% Tween-20 and 0.5% PPR negative goat serum) was added to all the wells of the ELISA plate. This was followed by the addition of the blocking buffer. The ELISA plate was kept in the incubator for 1 h at 37°C and thereafter the plate was washed. 100 μl of the primary antibody was added. This was followed by 1 h incubation at 37°C and washing. Diluted (1:1000) anti-mouse conjugate (100 μl) was added to all the wells of the plate. The ELISA plate was again kept in the incubator on an orbital shaker at 37°C for incubation up to 1 h. The discarding and washing procedures were repeated. 100 μl of OPD(o-phenylenediamine dihydrochloride) substrate solution was prepared. The solution was added to each well of the plate and the plate was kept in the incubator at 37°C for 10-20 minutes without shaking. The reaction was deemed to have completed when color developed in the positive reference (C+) wells. Thereafter, 100 μl stopping solution was dispensed in each well, and using an ELISA plate reader (Multiskan plus, LabSystem), the ELISA plate was read at 490 nm wavelength.

### RT-PCR

RNA isolation was carried out using the Trizol method. Nanodrop (Thermo Scientific™) was used to measure the quantity of RNA and to assess protein and salt. The RNA integrity was assessed by running the RNA samples in TapeStation (Agilent). cDNA was prepared through PrimeScript™ 1st strand cDNA Synthesis Kit (cat #6110A). Diagnostic PCR for the N gene was carried out with the help of published primers [26]. 10 μl of the reaction mixture was prepared with 5 μl of SapphireAmp Fast PCR Master Mix (Takara #RR350B), 1 μl of each of forward and reverse primers (final concentration 0.2 μM), 1 μl of 100ng template cDNA, and 2 μl of Nuclease Free Water (NFW). PCR was carried out with an initial denaturation at 95°C for 5 min followed by 35 cycles of denaturation at 95°C for 30 secs, annealing at 60°C for 30 secs, and renaturation at 72°C for 30 secs, with a final extension step at 72°C for 5 min. The PCR product was visualized on 1.5% agarose gel.

### m^6^A-sequencing

m^6^A-seq was carried out as previously described [14] with minor chamnges .RNA of around 200 μg was fragmented using RNA fragmentation buffer (Merck #CS220011) at 94°C for 5 minutes. The fragmentation reaction was stopped by the addition of EDTA (Merck #CS203175). The purified fragmented RNA was precipitated by using Sodium Acetate (Sigma #S2889-250G), Glycogen, and 100% ethanol. The size of the fragments was assessed through Tapestation. Around 100ng of the RNA was kept as input control and stored at -80°C whereas the remaining amount was subjected to immunoprecipitation using m^6^A specific antibody (NEB #E1610S). Initially, 30 μg of Protein G magnetic beads (NEB #S1430) was washed with 250 μl of Immunoprecipitation Reaction Buffer (150 mM NaCl, 10 mM Tris-HCl, pH 7.5, 0.1% NP-40 in NFW) and using a magnetic rack (NEB #S1506),the buffer was decanted after equilibration. The step was repeated once. Then, 10 μl of m^6^A antibody along with 200 ul Immunoprecipitation Reaction Buffer (IP) was added to the beads and incubated with head-over-tail rotation for 2 hours at 4°C. This was followed by the removal of the buffer by placing the microtube in a magnetic rack and three times washing with IP buffer.200 μl IP buffer wasalong with fragmented RNA and 10 μl RNAse Inhibitor (Merck #CS216138) was then added to the antibody coupled to beads. The reaction mixture was incubated overnight at 4°C with head-over-tail rotation. This was followed by decanting off the buffer through the magnetic rack. The sample mixture was washed twice with 500 μl IP buffer, twice with low salt wash buffer (50 mM NaCl, 10 mM Tris-HCl, pH 7.5, 0.1% NP-40 in NFW), twice with high salt wash buffer (500 mM NaCl, 10 mM Tris-HCl, pH 7.5, 0.1% NP-40 in NFW), and once again with IP buffer. For elution of m^6^A immunoprecipitated RNA, 100 μl of elution buffer [20mM m^6^A Salt (Merck #CS220007) and IP Buffer] was added to the beads and incubated at 4°C with head-over-tail rotation for 1 hour. Thereafter, the elution buffer was transferred into a 1.5 ml microtube. The steps were repeated with another 100 ml volume of elution buffer. RNA was precipitated from a total of 200 μl of elution volume using ethanol, sodium acetate, and glycogen. RNA sequencing of the eluted RNA, as well as the input RNA, was carried out thereafter.

### m^6^A sequencing analysis

The sequencing reads were initially screened for their quality through FastQC software [27]. Then, paired-end sequencing data of each sample was processed for the removal of adapters, low-quality reads, and singleton reads through Trimmomatic[28].The reference genome and the corresponding gtf file of *Capra hircus*[29] were retrieved from Ensembl. Thereafter, paired-end data of each sample was mapped to the *Capra hircus* genome using HISAT2 [30]. Samtools was used to convert sam files to bam format and to sort the bam files [31]. The PCR duplicates present in the bam files were removed through Picard tools. MeTDiff was used to find the m^6^A peaks that vary between the infected and non-infected samples with parameters: WINDOW_WIDTH=50, SLIDING_STEP=50, FRAGMENT_LENGTH=100, PEAK_CUTOFF_FDR=0.05, FOLD_ENRICHMENT=1 and DIFF_PEAK_CUTOFF_FDR=0.05 [32]. ChipSeekeer[33] was used to annotate m^6^A peaks from the output bed files from MetDiff. Customized code was used to count the number of altered m^6^A per gene and per chromosome. ggplot2 was used to plot figures.

### RNA -sequencing analysis

The filtered sequencing data of the “INPUT RNA” of each sample was used for RNA-sequencing analysis. The reference genome was prepared and indexed by RNA-Seq by Expectation-Maximization (RSEM) by rsem-prepare-reference command [34]. Further, the clean reads obtained from filtering of raw data were aligned to the indexed reference genome by Bowtie2[35]to estimate gene abundance in counts by rsem-calculate-expression command. To compare the gene expression levels between PPRV infective lung and PPRV negative lung, the aligned reads were used to generate a data matrix by ‘rsem-generate-data-matrix’ command and differential gene expression analysis was carried out by edgeR package [36]. The Ensembl IDs of the differentially expressed genes (DEGs) were converted to the respective gene ID by biomart[37], and the Reactome database was used for functional analysis [38].

### Real-time PCR

To validate the expression of the m^6^A related enzymes, qRT-PCR was done. HMBS (Hydroxymethylbilane Synthase) was taken as the internal control as it was found to be the best suitable endogenous control in gene expression studies of PPRV infection in tissues[39]. Each of the samples was run in triplicates and the relative expression of each gene was calculated using the 2^− ΔΔCT^ method with control as the calibrator[40]. The student’s t-test was done in JMP9 (SAS Institute Inc., Cary, USA) to test the significance of the difference. Differences between groups were considered significant at P ≤ 0.05.

## Results

### Confirmation of the PPRV infection

PPRV specific N gene amplicon size of 351 bp confirmed the presence of PPRV infection in the lung of PPRV infected goats and the absence of PPRV in the lung of healthy goats. Sandwich ELISA revealed the absence of PPRV antigen in the tissue samples of apparently healthy goats and the presence of the PPRV antigen in the lung tissues of PPRV infected goats **(Supplementary File 1)**.

### Alteration in m^6^A Peaks in RNA of Lung of Goats under PPRV Infection

To find m^6^A peaks that change under PPRV infection, m^6^A seq data of infected lungs PPRV infected goat was compared with the lung of PPRV negative goat using MetDiff. m^6^A-seq analysis revealed 1289 m^6^A peaks to be significantly altered [FDR < 0.05, |FC| > 0.5] in PPRV infected lung in comparison to normal lung, out of which 975 m^6^A peaks were hypomethylated [FDR < 0.05, FC < –0.5] and 314 peaks were hypermethylated [FDR < 0.05, FC > +0.5] **(Supplementary File 2, Figure 2A, 2B)**. As multiple m^6^A peaks can be found in a single mRNA [13], expectedly, there can be alteration in m^6^A peaks at more than one site within the same mRNA. A total of 1289 altered m^6^A peaks were found to be distributed in the transcripts of 868 genes, out of which 643 genes contained only one altered m^6^A peak, whereas 134 and 50 genes were found to change two and three positions respectively **(Figure 2C, Supplementary File 3)**. We then investigated the distribution of the m^6^A sites in the mRNA. Metaplot analysis revealed a higher distribution of m^6^A peaks near the stop codon in the case of the control sample while a shift away from the stop codon was observed under PPRV infection. UTR regions remained unperturbed **(Figure 2D, 2E)**.Chipseeker results revealed an equitable loss of m^6^A peaks from UTRs and CDS (Relative loss of m^6^A peaks = no. of hypomethylation/ total number of m^6^A peaks) **(Figure 3A, 3B)**. Overall, 32 genes were found to undergo hypomethylation as well as hypermethylation **(Figure 3C)**. The distribution of differentially methylated m^6^A containing genes across different chromosomes indicated relatively higher peaks in chromosomes 5, 13, and 19 **(Figure 3D)**.

**2.**
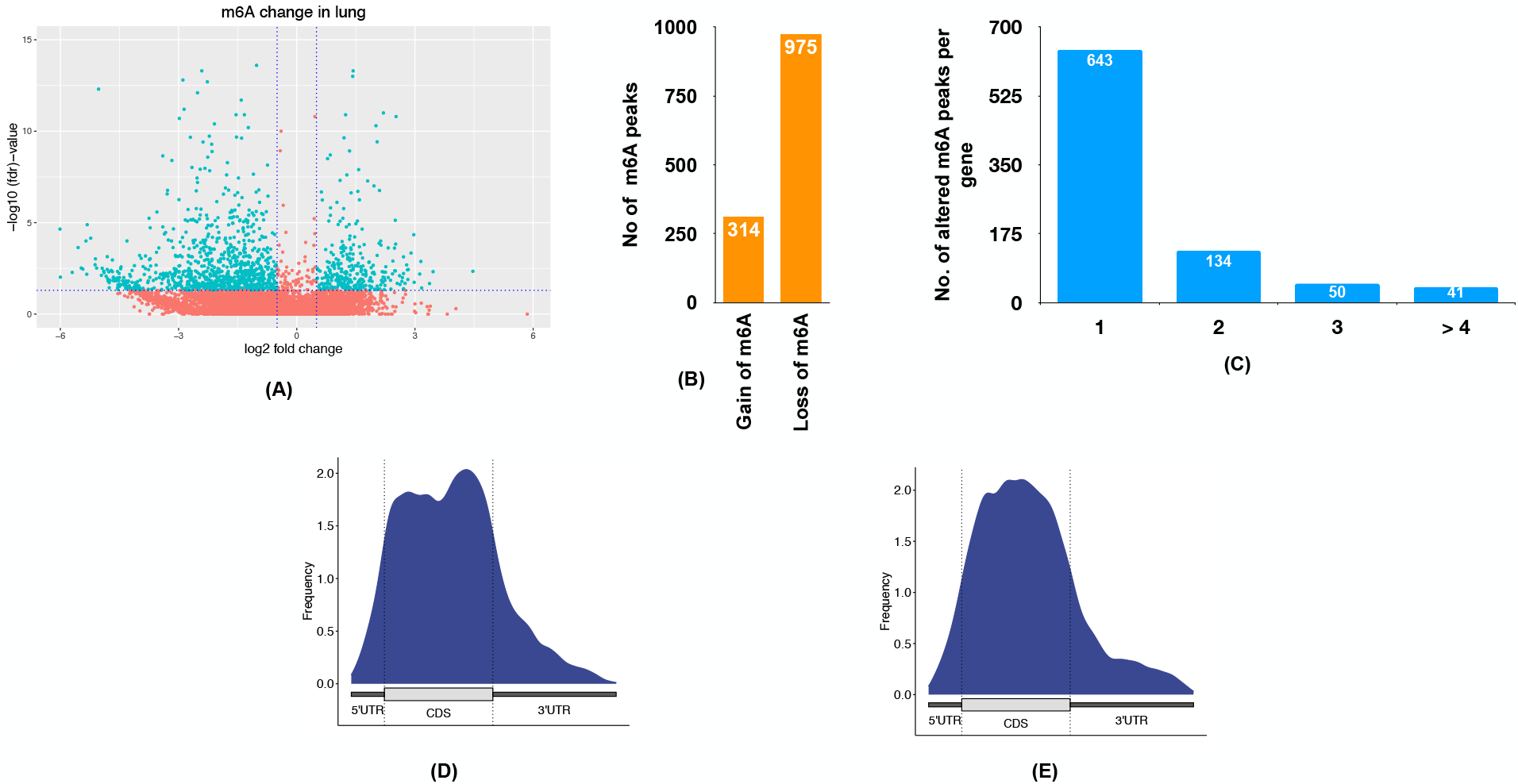
Change in m^6^A peaks following PPRV infection. **(A)** Volcano Plot of differentially methylated genes in lung PPRV infection. The X-axis is the log2ratio of methylation level between the infected and control lung. The Y-axis is the differential FDR based on a negative log scale. The cyan dots represent the significant differential m^6^A peaks, based on FDR< 0.05 and |FC| > 0.5. (B) Bar diagram showing the number of hypomethylated and hypermethylated m^6^A peaks (C) The distribution of differential m^6^A peaks per gene (D) Metaplot of m^6^A peaks in control lung (E) Metaplot of m^6^A peaks in PPRV infected lung

**3.**
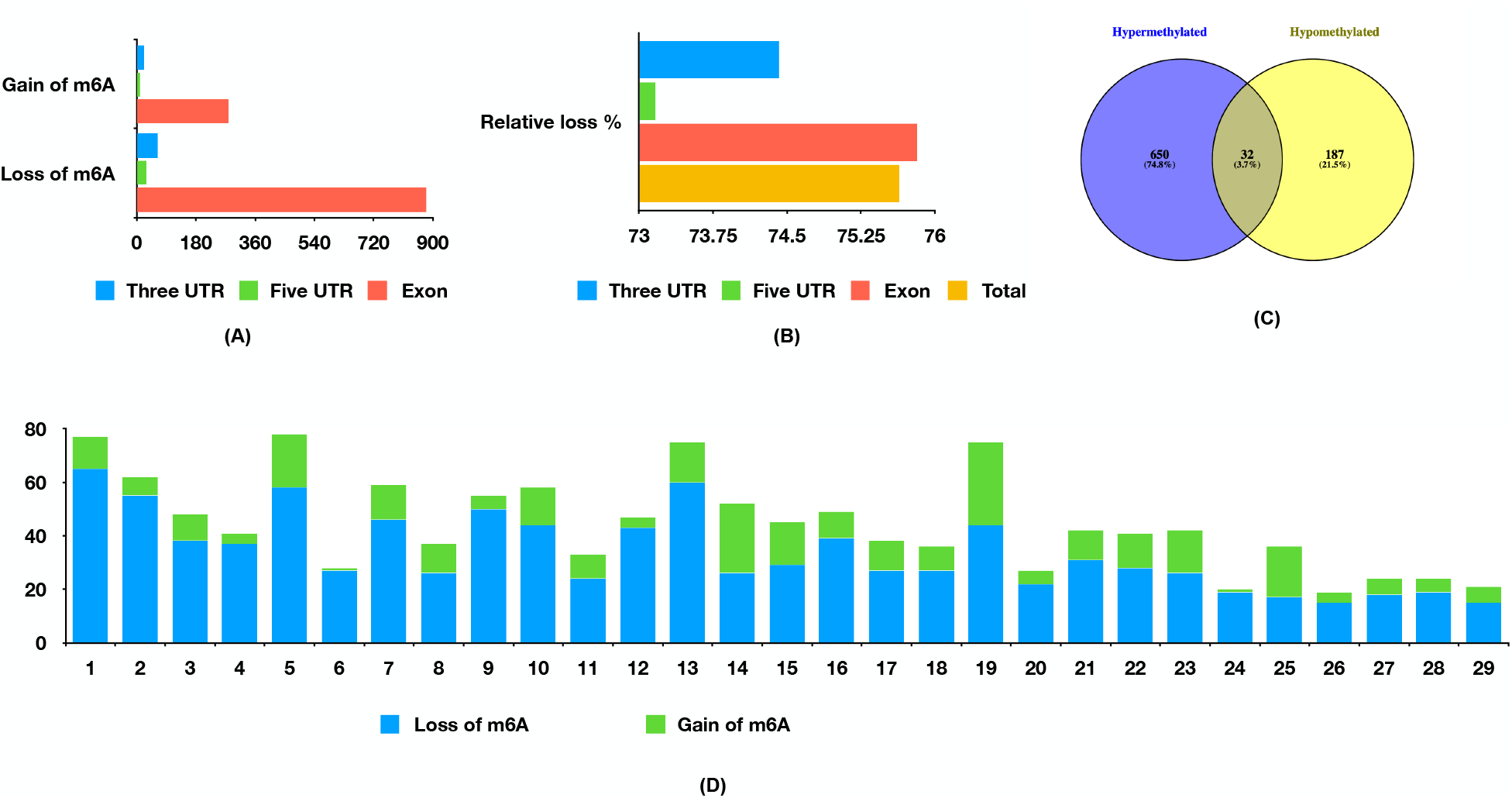
(A) The number of altered m^6^A peaks in CDS and UTR regions (B) Relative loss of m^6^A peaks (= no. of hypomethylation/ total number of altered m^6^A peaks *100) from UTR and CDS region (C) Venn diagram showing the genes that are hypermethylated at a certain position(s) and hypomethylated at other position(s) (D) Distribution of hypomethylated and hypermethylated m^6^A peaks in different chromosomes.

### Functional analysis of differentially m^6^A containing genes

Through Reactome, we carried out functional analysis of the genes that had undergone only hypomethylation, only hypermethylation, and both hypermethylation as well as hypomethylation. We found out that hypermethylated genes are enriched for pathways that include PKMTs methylate histone lysines, synthesis of PIPs at the plasma membrane, RUNX1 regulates genes involved in megakaryocyte differentiation and platelet function, Signaling by Interleukins, and TCF dependent signaling in response to WNT **(Figure 4A)**. Hypomethylated genes were enriched in Interleukin-4 and Interleukin-13 signaling and various processes associated with extracellular matrix organization **(Figure 4C)**. The genes that undergo both hypomethylation as well as hypermethylation were found enriched in various RNA metabolic processes like mRNA editing, miRNA biogenesis, and repression and transcription regulation **(Figure 4B)**.

**4.**
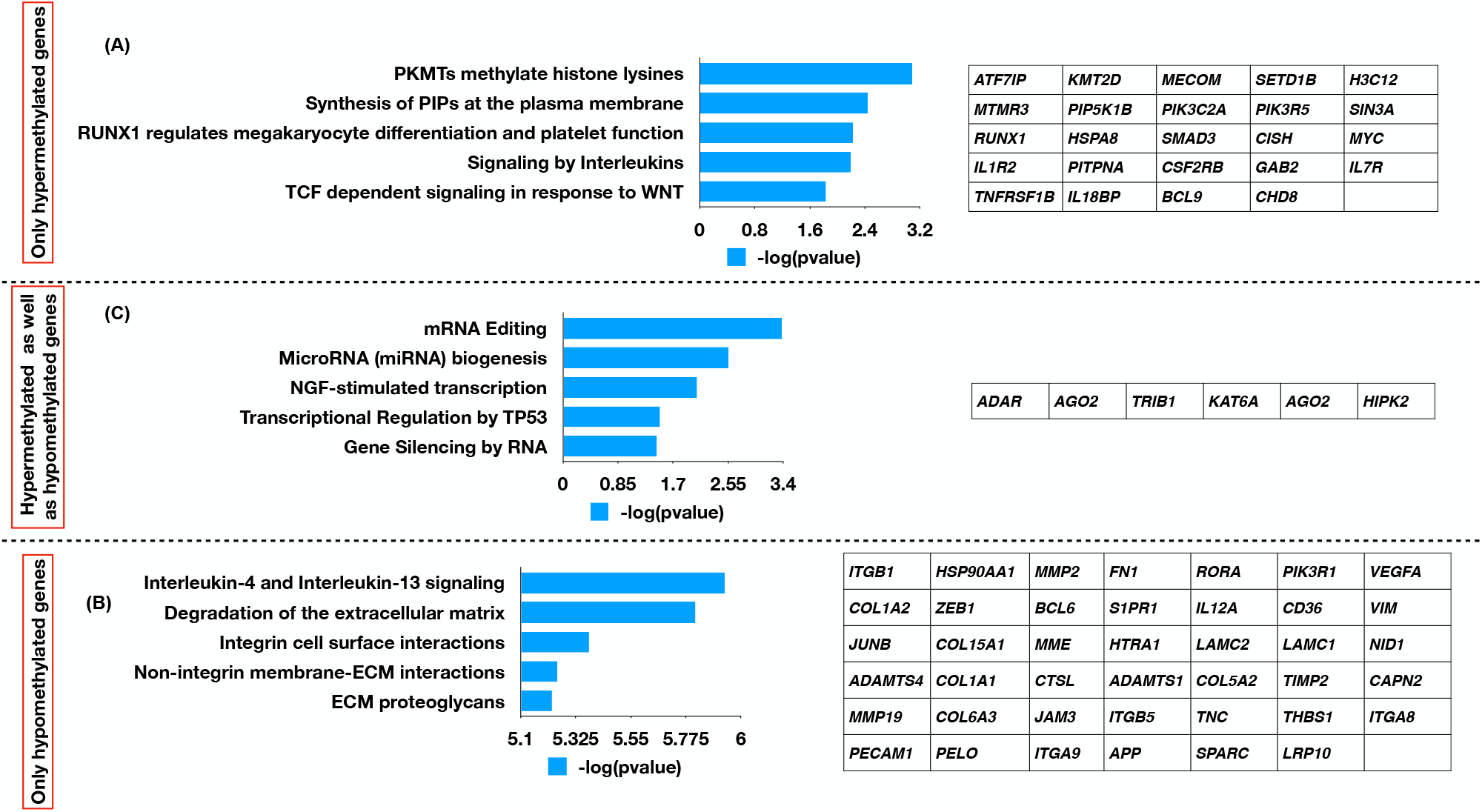
The functional analysis of genes with (A) m^6^A hypermethylated (C) m^6^A hypomethylated peaks (B) having both hypermethylated and hypomethylated peaks. The significance of the process is based on p-value as indicated by –log10 (P) on X-axis.

### Functional analysis of differentially expressed genes

The input sequencing reads (input RNA) of each sample were used for RNA sequencing analysis. The differential expression of genes in the lungs of PPRV infected animals in comparison to PPRV negative animals using edgeR revealed dysregulation of 5743 genes following PPRV infection, out of which 2547 were upregulated and 3196 were downregulated **(Supplementary File 4, Figure 5A, 5B)**. The upregulated genes were highly enriched in processes like neutrophil degranulation, rRNA modification in the nucleus and cytosol, Unfolded Protein Response (UPR), cellular response to heat stress, antiviral mechanism by IFN-stimulated genes, cytokine signaling in immune system, and signaling by Interleukins **(Figure 5C)** while downregulated genes were enriched in eukaryotic translation elongation, eukaryotic translation termination, viral mRNA Translation, nonsense-mediated decay, and extracellular matrix organization **(Figure 5D)**. We then looked into the genes related to the significant processes that were found to be enriched by the upregulated genes and downregulated genes (**Figure 5E)**.

**5.**
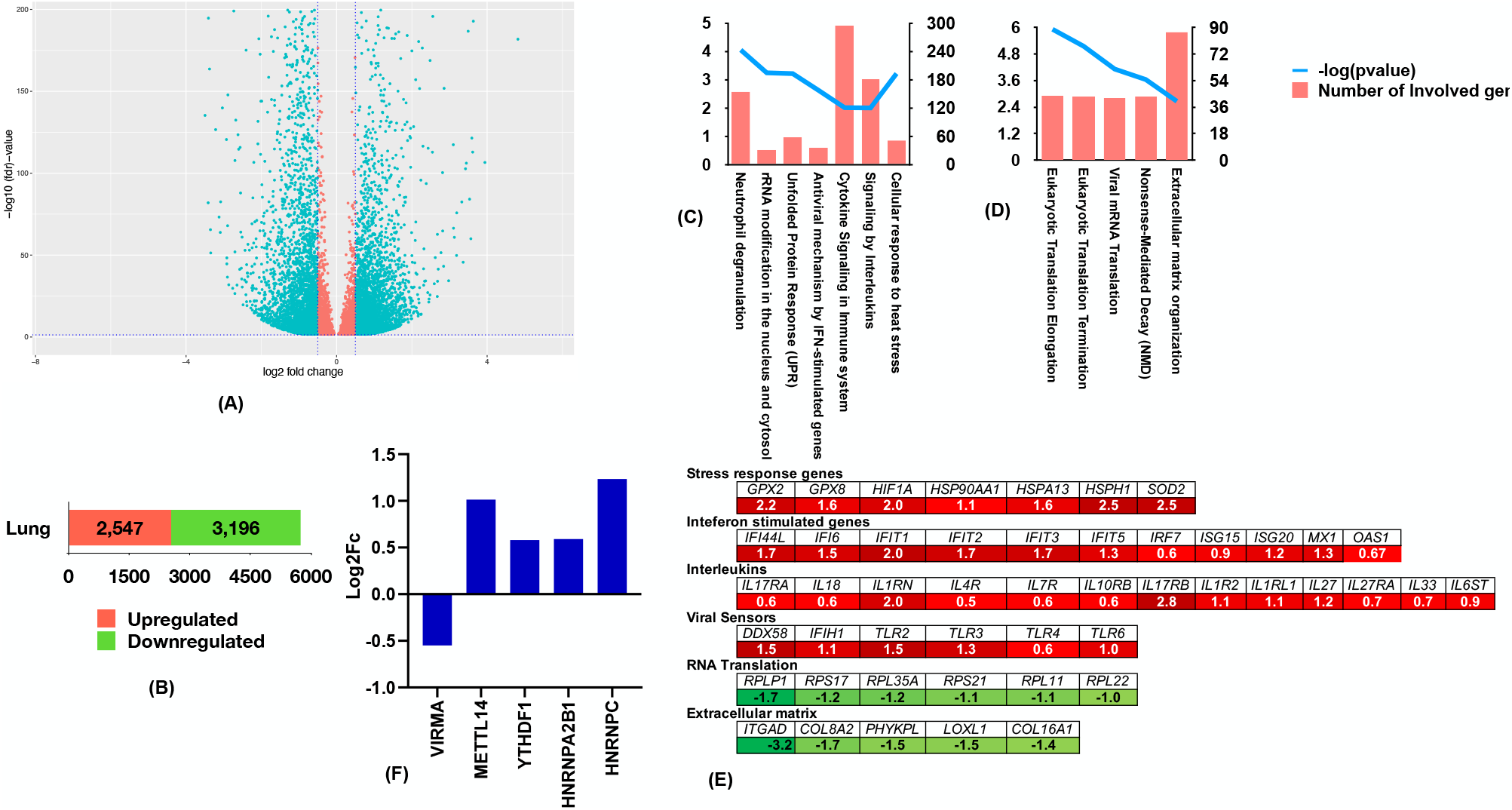
Change in expression of genes following PPRV infection. (A) Volcano Plot of differentially expressed genes in the spleen following PPRV infection. The X-axis is the log2 ratio of expression level between the infected and control lung. The Y-axis is the differential FDR based on negative log scale. The cyan dots represent significant DEGs based on FDR < 0.05 and |FC| > 0.5 (B) The number of upregulated and downregulated m^6^A peaks (C) Functional analysis of upregulated genes (D) Functional analysis of downregulated genes. The significance of the process in figures (C) and (D) is based on p-value as indicated by –log10 (P) on the left Y-axis. The right Y-axis in Figures (C) and (D) indicate the number of genes involved in the processes. (E) Alteration in expression of enzymes related to m6A regulatory machinery.(F) Log_2_FC of genes involved in the different processes associated with DEGs. The red color indicates upregulated genes and the green color indicates downregulated genes

### Expression of Immune genes increased following PPRV infection

In earlier studies of host-PPRV interaction, we have reported higher expression of genes encoding different molecule species such as Interferon-Stimulated Genes (ISGs), interleukins and viral sensors involved in innate immunity following PPRV infection [7, 41]. Among the immune-related processes enriched by the upregulated genes, we found that upregulated genes involved in these processes are ISGs like *IFI44L, IFI6, IFIT1, IFIT2, IFIT3, IFIT5, IRF7, ISG15, ISG20*, and *MX1*; interleukin and interleukin receptors such as *IL17RA, IL18, IL1RN, IL4R, IL7R, IL10RB, IL17RB, IL1R2, IL1RL1, IL27, IL27RA, IL33*, and *IL6ST*, and viral sensors like RIG-I (*DDX58*), MDA5 (*IFIH1*) and *TLR2, TLR3, TLR4*, and *TLR6* **(Figure 5E)**.

### Expression of stress related genes increased following PPRV infection

GPX and SOD are two components of the antioxidant triad that forms the first line of defence against reactive oxygen species[42-44]. The expression of GPX and SOD was found to increase following PPRV infection. Further, mRNAs encoding heat shock proteins such as heat shock protein HSP 90-alpha *(HSP90AA1*), heat shock 70 kDa protein 13 *(HSPA13)*,heat shock protein 105 kDa (*HSPH1*) were found to be elevated **(Figure 5E)**.

### Genes involved in extracellular matrix and RNA translation were downregulated under PPRV infection

Ribosomal proteins are vital for the translation of RNA. Few of the downregulated genes involved in the processes such as eukaryotic translation elongation, eukaryotic translation termination, viral mRNA included *RPLP1, RPS17, RPL35A, RPS21, RPL11*, and *RPL22*. Besides, certain downregulated genes involved in the extracellular matrix organization included Integrin alpha-D (*ITGAD)* and the *COL8A2, PHYKPL, LOXL1*, and *COL16A1* which are required for collagen synthesis **(Figure 5E)**.

We also screened the DEGs for the expression of known m^6^A writers, readers, and erasers. We found m^6^A reader *HNRNPC* and the core component of methyltransferase complex *METTL14* to be highly upregulated than the m^6^A readers – *HNRNPA2B1* and *YTHDF1*. We also found the accessory component of METTL3/14 complex – *VIRMA* to be downregulated **(Figure 5F)**. The expression of VIRMA, METTL14, and YTHDF1 was confirmed through qPCR **(Supplementary File 5)**.

### Correlation of DEGs with differentially methylated genes

m^6^A modification influences the stability of the m^6^A containing RNAs – it has been found to facilitate RNA degradation and stabilize the RNA as well and the process is contingent upon the location of m^6^A on the transcript and the involvement of m^6^A readers [45-48]. We, therefore, correlated the expression of DEGs with that of genes containing differential m^6^A peaks. Of the 843 differentially m^6^A containing genes, 282 genes were found to be differentially expressed. The 282 DEGs were categorized into 4 types of combinations 1) hypermethylated and upregulated genes 2) hypermethylated and downregulated 3) hypomethylated and upregulated genes 4) hypomethylated and downregulated genes. It was found that out of 282 DEGs, 103 downregulated genes were hypomethylated and 27 were hypermethylated. Also, 25 upregulated gene was found hypermethylated and 116 were hypomethylated. Further, the proportion of DEGs that underwent a change in m^6^A methylation at a single position was found far higher than the DEGs in which more than one m^6^A methylation site changed following PPRV infection **(Figure 6A, 6B)**. Functional analysis carried revealed that the 282 genes are significantly enriched in signaling by Interleukins, extracellular matrix organization, cytokine signaling in Immune system, signaling by receptor tyrosine kinases, and Toll-like Receptor Cascades **(Figure 6C)**. Among the interleukin family, a gain of m^6^A methylation was found in upregulated mRNAsofinterleukin receptors *– IL1R2* and *IL7R*, and the loss of m^6^A methylation in upregulated *IL17RA* mRNA. Besides, loss of m^6^A modification at three positions was found in downregulated mRNA of subunit A component of Complement C1q (*C1QA*). The only ISG in which there was a gain of m^6^A was *OAS1*. Moreover, among the downregulated genes, genes involved in extracellular matrix organization – COL15A1, ITGA8, and LOXL1 were found to undergo loss of m^6^A following PPRV infection. Moreover, loss of m^6^A was also found in mRNA of upregulated HSP90 and HSP70 **(Figure 6D)**.

**6.**
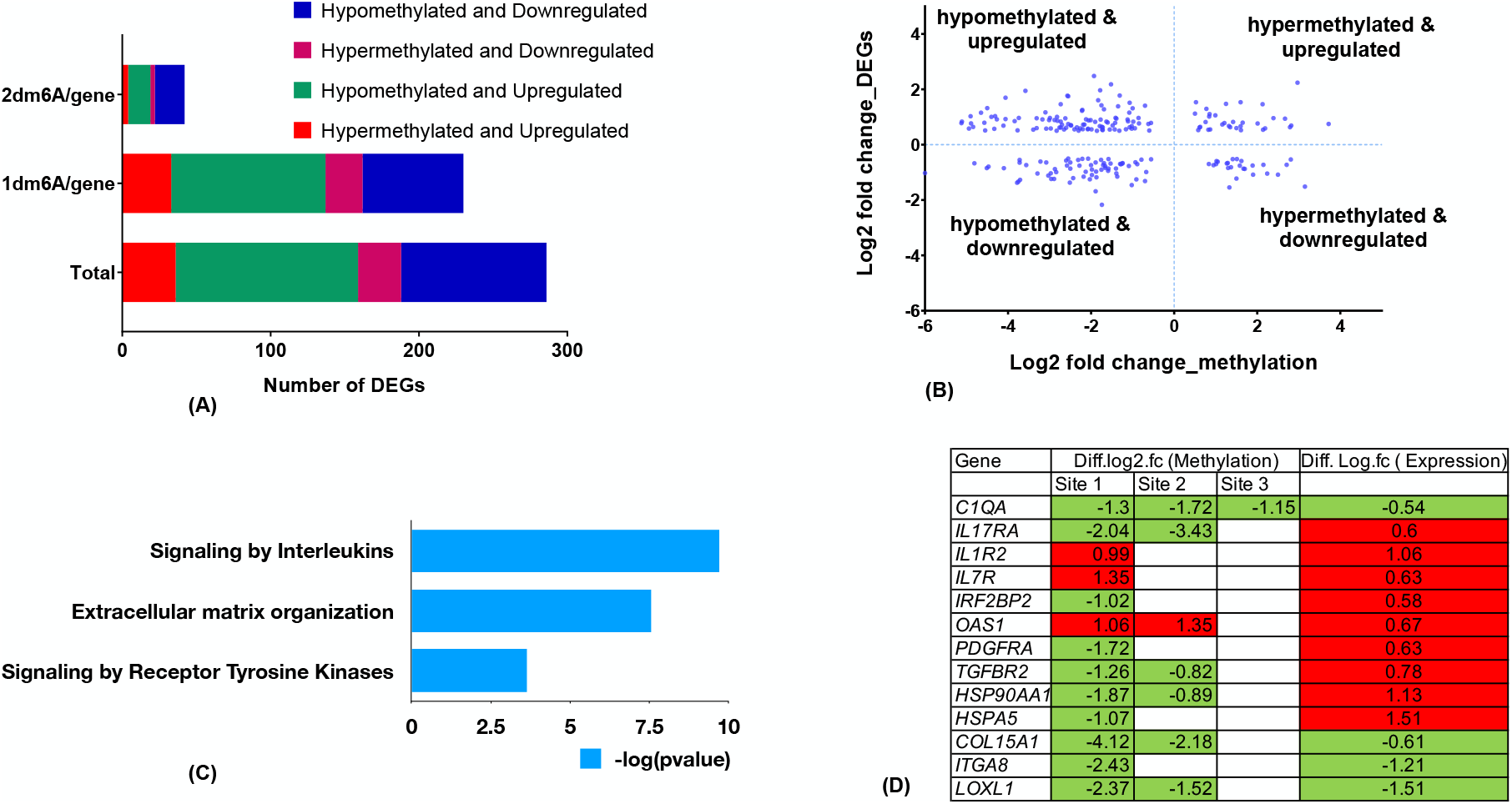
Correlation between differential m^6^A-containing peaks and differentially expressed genes: **(A)** The changes in m^6^A peaks of differentially expressed genes: total, at one position (1dm^6^A), and at two positions (2dm^6^A).The combinations shown include Hypermethylated and upregulated genes; Hypermethylated and downregulated; Hypomethylated and upregulated genes, and Hypomethylated and downregulated genes in the lung following PPRV infection. (B) Quadrant plots of differentially expressed genes in which there was alteration at one site. The X-axis is the log2 ratio of methylation level between the infected and control tissue. The Y-axis is the log2 ratio of expression level between the infected and control tissue (C) Functional analysis of the DEGs which contain differential m^6^A peaks (D) Log2FC values of change in methylation and expression of some important genes

## Discussion

PPR is a contagious disease of sheep and goats. Because of wide geographical distribution with high mortality and morbidity, PPR is an economically important disease for small ruminant farming. Recently, high throughput techniques have been used for a better understanding of the molecular pathogenesis of PPRV. The immune mechanisms activated by the attenuated vaccine strain and virulent strains of PPRV indicate similar expression patterns [5, 7, 41, 49]. However, differences in the degree of expression have been observed that pointed to the role of other regulatory mechanisms in PPRV infections.

We divulged one of the most abundant and functionally recognized RNA modifications – m^6^A modification of host mRNA under PPRV infection. Goats were initially experimentally infected with the Izatnagar/94 strain[23] of the virus and sacrificed following the development of the clinical presentation of PPRV infection on the 7th-day post-infection. Bilateral mucopurulent discharge, coughing, fetid diarrhea, and general weakness were observed [24]. Among all the transcripts of PPRV, N mRNA is most transcribed and is used to confirm PPRV infection [7-9]. Similarly, in the present study, we confirmed experimental PPRV infection in goats using N gene expression and sandwich ELISA.

We carried out m^6^A sequencing analysis of the infected samples as well as a noninfected sample (n=2) to identify the m^6^A peaks that change following PPRV infection. 1289 m^6^A peaks were found to be significantly altered in PPRV infected lungs in comparison to the control lungs. The results indicated a gain of m^6^A as well as loss of m^6^A taking place on mRNAs following PPRV infection. The significant alteration in the m^6^A landscape of host mRNA has been reported in other m^6^A studies of different host-viral interactions[15, 18, 50].

Our analysis revealed that the altered m^6^A peaks represent transcripts of 868 genes. We found that certain mRNAs undergo m^6^A change at more than one position, however, their proportion was lower as compared to those that change at a single point. In the case of the control lung, the m^6^A peaks were found within the UTR and CDS regions but highly enriched near the stop codons as has been reported in other mammals [14]. Earlier studies indicate that the change in the distribution of m^6^A peaks can be virus-specific as well as cell-specific. The distribution of m^6^A on host mRNA does not change following Zika virus, dengue virus, hepatitis C virus, and West Nile virus [50]. Among the four different cell types infected with KSHV, a decrease within the 5’ UTR and a corresponding increase within the 3’ UTR was found in three cell types[18]. However, no such redistribution of m^6^A peaks of host mRNA after PPRV infection was found.

We also found that expression of m^6^A writer METTL14 and an m^6^A reader HNRNPC was elevated at the transcriptome level following PPRV infection. However, the expression of m^6^A erasers FTO and ALKBH5 was found to be unperturbed. Thus, the cause-effect relation of the increase in the number of hypomethylated peaks with the expression of m^6^A regulatory enzymes could not be linked in PPRV infected lungs. Recently transcription factors, histone modification, and RNA binding proteins have been found to regulate the recruitment of methyltransferase complex on target mRNAs [51-53]. Moreover, a decrease in transcriptional rate has been shown to increase the number of m^6^A sites [54]. The intricate influence of various accessory factors could also influence alteration in m^6^A peaks following PPRV infection.

Functional analysis of upregulated genes showed enrichment in IFN-stimulated genes, cytokine signaling in immune system, and signaling by Interleukins. Notable upregulated genes involved in these are ISGs, interleukin and interleukin receptors, and viral sensors. ISGs – ISG15, ISG20, IRF7, IFIT1, IFIT3, IFIT5, MX1, IFI6, and OAS1 whose antiviral activity has been reported in various RNA virus infections [55], have also been found to be elevated in lymphocytes isolated from virulent PPRV infected goat [7] and various PBMCs subsets isolated from Sungri 96/PPRV vaccine goats [41].

The expression pattern of viral sensors RIG-I and MDA5 which are required for stimulation of ISGs[56] is similar to virulent PPRV infected lymphocytes [7]andPBMCs from PPRV vaccinated animals[41]. Moreover, upregulated genes were also found enriched in stress response genes. The molecules involved are antioxidants – SOD2 and GPX2 which alleviate the virus-induced oxidative injury [57]. Ribosomal proteins like RPLP1, RPS17, RPL11, and RPL22 which have a positive influence on viral propagation are downregulated [58].

We also found that hypermethylated transcripts were enriched in processes that are associated with chromosomal organization, synthesis of PIPs at the plasma membrane, megakaryocyte differentiation and platelet function, Interleukin signaling, and WNT signaling. The hypomethylated genes were associated specifically with Interleukin-4 and Interleukin-13 signaling and various processes associated with extracellular matrix organization. The alteration in m^6^A under PPRV infection is not just limited to transcripts of specific processes or signaling but ranges from different innate immune processes to the processes related to cell proliferation and extracellular organization. m^6^A regulates RNA stability, RNA translation, splicing, and export from the nucleus [12]. To analyze the effect of altered m^6^A modification on gene expression, we correlated the RNA sequencing data with m^6^A-seq data. Both loss and gain of m^6^A modification were found in upregulated as well as downregulated genes, which is in coherence with the studies about stabilizing as well as the destabilizing effect of m^6^A [45-47]. We found that the proportion of hypomethylated downregulated and hypomethylated upregulated is very high than the proportion of hypermethylated downregulated and hypermethylated upregulated. No change in m^6^A was found in upregulated mRNAs of ISGs, interleukins, and viral sensors. However, upregulated genes of interleukin receptors such as IL1R2, IL7R and IL17RA were found where an alteration in m^6^A sites was found. m^6^A modification seems to play an important role in regulating interleukin expression and ISG expression during PPRV.

## Conclusion

Through m^6^A-seq analysis of lung of goat, m^6^A peaks were found to be significantly altered following PPRV infection. The m^6^A hypomethylated genes were found enriched in Interleukin-4 and Interleukin-13 signaling. A fraction of differentially expressed genes with alteration of m^6^A modification was found to belong to cytokine signaling pathway. The involvement of m^6^A modification adds another connecting link between the PPRV infection and the host.

## Supporting information

Supplementary Fille 3

Supplementary Fille 4

Supplementary Fille 5

## Data Availability

All raw sequencing data generated in this study are available in GEO, NCBI under accession number GSE190718.

## Author contributions

**Raja Ishaq Nabi Khan:** Investigation, Writing - Original Draft, Formal analysis, Visualization; **Manas Ranjan Praharaj:** Investigation, Writing - Review & Editing, Software; **Waseem Akram Malla:** Investigation, Writing - Review & Editing, Software; **NeelimaHosamani:** Investigation, Visualization; **Shikha Saxena:** Investigation, Visualization; **Bina Mishra:** Writing - Review & Editing; **Kaushal Kishor Rajak:** Investigation, Methodology; **MuthuchelvanDhanavelu:** Investigation, Methodology; **Ashok Kumar Tiwari:** Resources, Conceptualization, Methodology; **B P Mishra:** Resources, Conceptualization, Methodology, Supervision, Funding acquisition; **Basavaraj Sajjanar:** Resources, Conceptualization, Methodology, Funding acquisition; **Ravi Kumar Gandham:** Resources, Conceptualization, Methodology, Funding acquisition

## Acknowledgements

We thank the Department of Biotechnology, Government of India for providing fellowship and contingency for Raja Ishaq Nabi Khan (DBT Fellow No. DBT/2017/IVRI/768).

## References

1. Muthuchelvan, D., et al., Peste-des-petits-ruminants: An Indian perspective. Adv. Anim. Vet. Sci, 2015. 3(8): p. 422–429.

2. Kumar, P., et al., Pathological and immunohistochemical study of experimental peste des petits ruminants virus infection in goats. Journal of Veterinary Medicine, Series B, 2004. 51(4): p. 153–159.

3. Banyard, A.C., et al., Global distribution of peste des petits ruminants virus and prospects for improved diagnosis and control. Journal of general virology, 2010. 91(12): p. 2885–2897.

4. Singh, R., U. De, and K. Pandey, Virological and antigenic characterization of two peste des petits ruminants (PPR) vaccine viruses of Indian origin. Comparative Immunology, Microbiology and Infectious Diseases, 2010. 33(4): p. 343–353.

5. Manjunath, S., et al., Genomic analysis of host–Peste des petits ruminants vaccine viral transcriptome uncovers transcription factors modulating immune regulatory pathways. Veterinary research, 2015. 46(1): p. 1–15.

6. Tirumurugaan, K.G., et al., RNAseq Reveals the Contribution of Interferon Stimulated Genes to the Increased Host Defense and Decreased PPR Viral Replication in Cattle. Viruses, 2020. 12(4): p. 463.

7. Wani, S.A., et al., Contrasting gene expression profiles of monocytes and lymphocytes from Peste-des-petits-ruminants virus infected goats. Frontiers in Immunology, 2019. 10.

8. Khanduri, A., et al., Dysregulated miRNAome and proteome of PPRV infected goat PBMCs reveal a coordinated immune response. Frontiers in immunology, 2018. 9: p. 2631.

9. Pandey, A., et al., Modulation of host miRNAs transcriptome in lung and spleen of Peste des Petits ruminants virus infected sheep and goats. Frontiers in microbiology, 2017. 8: p. 1146.

10. Boccaletto, P., et al., MODOMICS: a database of RNA modification pathways. 2017 update. Nucleic acids research, 2018. 46(D1): p. D303–D307.

11. Khan, R.I.N. and W.A. Malla, m(6)A modification of RNA and its role in cancer, with a special focus on lung cancer. Genomics, 2021. 113(4): p. 2860–2869.

12. He, P.C. and C. He, m6A RNA methylation: from mechanisms to therapeutic potential. The EMBO Journal, 2021. 40(3): p. e105977.

13. Meyer, K.D., et al., Comprehensive analysis of mRNA methylation reveals enrichment in 3’ UTRs and near stop codons. Cell, 2012. 149(7): p. 1635–46.

14. Dominissini, D., et al., Topology of the human and mouse m 6 A RNA methylomes revealed by m 6 A-seq. Nature, 2012. 485(7397): p. 201–206.

15. Lichinchi, G., et al., Dynamics of human and viral RNA methylation during Zika virus infection. Cell Host & Microbe, 2016. 20(5): p. 666–673.

16. Cristinelli, S., et al., HIV modifies the m6A and m5C epitranscriptomic landscape of the host cell. bioRxiv, 2021: p. 2021.01. 04.425358.

17. Lichinchi, G., et al., Dynamics of the human and viral m 6 A RNA methylomes during HIV-1 infection of T cells. Nature microbiology, 2016. 1(4): p. 1–9.

18. Tan, B., et al., Viral and cellular N 6-methyladenosine and N 6, 20-O-dimethyladenosine epitranscriptomes in the KSHV life cycle. Nature microbiology, 2018. 3(1): p. 108–120.

19. Wani, S.A., et al., Expression kinetics of ISG15, IRF3, IFNγ, IL10, IL2 and IL4 genes vis-a-vis virus shedding, tissue tropism and antibody dynamics in PPRV vaccinated, challenged, infected sheep and goats. Microbial pathogenesis, 2018. 117: p. 206–218.

20. Chowdhury, E.H., et al., Natural peste des petits ruminants virus infection in Black Bengal goats: virological, pathological and immunohistochemical investigation. BMC Veterinary Research, 2014. 10(1): p. 1–10.

21. Singh, R.P., et al., Development of a monoclonal antibody based competitive-ELISA for detection and titration of antibodies to peste des petits ruminants (PPR) virus. Vet Microbiol, 2004. 98(1): p. 3–15.

22. Dhinakar Raj, G., K. Nachimuthu, and A. Mahalinga Nainar, A simplified objective method for quantification of peste des petits ruminants virus or neutralizing antibody. J Virol Methods, 2000. 89(1-2): p. 89–95.

23. Sahu, A.R., et al., Genome sequencing of an Indian peste des petits ruminants virus isolate, Izatnagar/94, and its implications for virus diversity, divergence and phylogeography. Archives of virology, 2017. 162(6): p. 1677–1693.

24. Pope, R.A., et al., Early events following experimental infection with peste-des-petits ruminants virus suggest immune cell targeting. PloS one, 2013. 8(2): p. e55830.

25. Singh, R.P., et al., A sandwich-ELISA for the diagnosis of Peste des petits ruminants (PPR) infection in small ruminants using anti-nucleocapsid protein monoclonal antibody. Arch Virol, 2004. 149(11): p. 2155–70.

26. Couacy-Hymann, E., et al., Rapid and sensitive detection of peste des petits ruminants virus by a polymerase chain reaction assay. Journal of virological methods, 2002. 100(1-2): p. 17–25.

27. Brown, J., M. Pirrung, and L.A. McCue, FQC Dashboard: integrates FastQC results into a web-based, interactive, and extensible FASTQ quality control tool. Bioinformatics, 2017. 33(19): p. 3137–3139.

28. Bolger, A.M., M. Lohse, and B. Usadel, Trimmomatic: a flexible trimmer for Illumina sequence data. Bioinformatics, 2014. 30(15): p. 2114–2120.

29. Bickhart, D.M., et al., Single-molecule sequencing and chromatin conformation capture enable de novo reference assembly of the domestic goat genome. Nature genetics, 2017. 49(4): p. 643–650.

30. Wen, G. A simple process of RNA-sequence analyses by Hisat2, Htseq and DESeq2. in Proceedings of the 2017 International Conference on Biomedical Engineering and Bioinformatics. 2017.

31. Li, H., et al., The sequence alignment/map format and SAMtools. Bioinformatics, 2009. 25(16): p. 2078–2079.

32. Cui, X., et al., MeTDiff: A Novel Differential RNA Methylation Analysis for MeRIP-Seq Data. IEEE/ACM Trans Comput Biol Bioinform, 2018. 15(2): p. 526–534.

33. Yu, G., L.-G. Wang, and Q.-Y. He, ChIPseeker: an R/Bioconductor package for ChIP peak annotation, comparison and visualization. Bioinformatics, 2015. 31(14): p. 2382–2383.

34. Pavet, V., et al., Towards novel paradigms for cancer therapy. Oncogene, 2011. 30(1): p. 1–20.

35. Langmead, B. and S.L. Salzberg, Fast gapped-read alignment with Bowtie 2. Nature methods, 2012. 9(4): p. 357–359.

36. Robinson, M.D., D.J. McCarthy, and G.K. Smyth, edgeR: a Bioconductor package for differential expression analysis of digital gene expression data. Bioinformatics, 2010. 26(1): p. 139–140.

37. Durinck, S., et al., Mapping identifiers for the integration of genomic datasets with the R/Bioconductor package biomaRt. Nature protocols, 2009. 4(8): p. 1184.

38. Fabregat, A., et al., The reactome pathway knowledgebase. Nucleic acids research, 2018. 46(D1): p. D649–D655.

39. Sahu, A.R., et al., Selection and validation of suitable reference genes for qPCR gene expression analysis in goats and sheep under Peste des petits ruminants virus (PPRV), lineage IV infection. Scientific reports, 2018. 8(1): p. 1–11.

40. Schmittgen, T.D. and K.J. Livak, Analyzing real-time PCR data by the comparative C T method. Nature protocols, 2008. 3(6): p. 1101.

41. Wani, S.A., et al., Systems Biology behind immunoprotection of both Sheep and Goats after Sungri/96 PPRV vaccination. Msystems, 2021. 6(2).

42. Lee, Y.S., et al., Dysregulation of adipose glutathione peroxidase 3 in obesity contributes to local and systemic oxidative stress. Molecular Endocrinology, 2008. 22(9): p. 2176–2189.

43. Bierl, C., et al., Determinants of human plasma glutathione peroxidase (GPx-3) expression. Journal of Biological Chemistry, 2004. 279(26): p. 26839–26845.

44. Ighodaro, O. and O. Akinloye, First line defence antioxidants-superoxide dismutase (SOD), catalase (CAT) and glutathione peroxidase (GPX): Their fundamental role in the entire antioxidant defence grid. Alexandria Journal of Medicine, 2018. 54(4): p. 287–293.

45. Shi, H., et al., YTHDF3 facilitates translation and decay of N 6-methyladenosine-modified RNA. Cell research, 2017. 27(3): p. 315–328.

46. Zhang, F., et al., Fragile X mental retardation protein modulates the stability of its m6A-marked messenger RNA targets. Human Molecular Genetics, 2018. 27(22): p. 3936–3950.

47. Chen, M., et al., RNA N6-methyladenosine methyltransferase-like 3 promotes liver cancer progression through YTHDF2-dependent posttranscriptional silencing of SOCS2. Hepatology, 2018. 67(6): p. 2254–2270.

48. Lasman, L., et al., Context-dependent functional compensation between Ythdf m6A reader proteins. Genes & development, 2020. 34(19-20): p. 1373–1391.

49. Manjunath, S., et al., Comparative and temporal transcriptome analysis of peste des petits ruminants virus infected goat peripheral blood mononuclear cells. Virus research, 2017. 229: p. 28–40.

50. Gokhale, N.S., et al., Altered m6A modification of specific cellular transcripts affects Flaviviridae infection. Molecular cell, 2020. 77(3): p. 542-555. e8.

51. Huang, H., et al., Histone H3 trimethylation at lysine 36 guides m(6)A RNA modification co-transcriptionally. Nature, 2019. 567(7748): p. 414–419.

52. Barbieri, I., et al., Promoter-bound METTL3 maintains myeloid leukaemia by m(6)A-dependent translation control. Nature, 2017. 552(7683): p. 126–131.

53. Fish, L., et al., Nuclear TARBP2 drives oncogenic dysregulation of RNA splicing and decay. Molecular cell, 2019. 75(5): p. 967-981. e9.

54. Slobodin, B., et al., Transcription impacts the efficiency of mRNA translation via co-transcriptional N6-adenosine methylation. Cell, 2017. 169(2): p. 326-337. e12.

55. Schneider, W.M., M.D. Chevillotte, and C.M. Rice, Interferon-stimulated genes: a complex web of host defenses. Annual review of immunology, 2014. 32: p. 513–545.

56. Park, A., A. Iwasaki, and microbe, Type I and type II. interferons–induction, signaling, evasion, and application to combat COVID-19. Cell Host & Microbe, 2020.

57. Camini, F.C., et al., Implications of oxidative stress on viral pathogenesis. Archives of Virology, 2017. 162(4): p. 907–917.

58. Belhadj Slimen, I., et al., Reactive oxygen species, heat stress and oxidative-induced mitochondrial damage. A review. International journal of hyperthermia, 2014. 30(7): p. 513–523.

