## Supplementary File 1 for "Changes in m^6^A RNA methylation of goat lung following PPRV infection"

### Slide 1
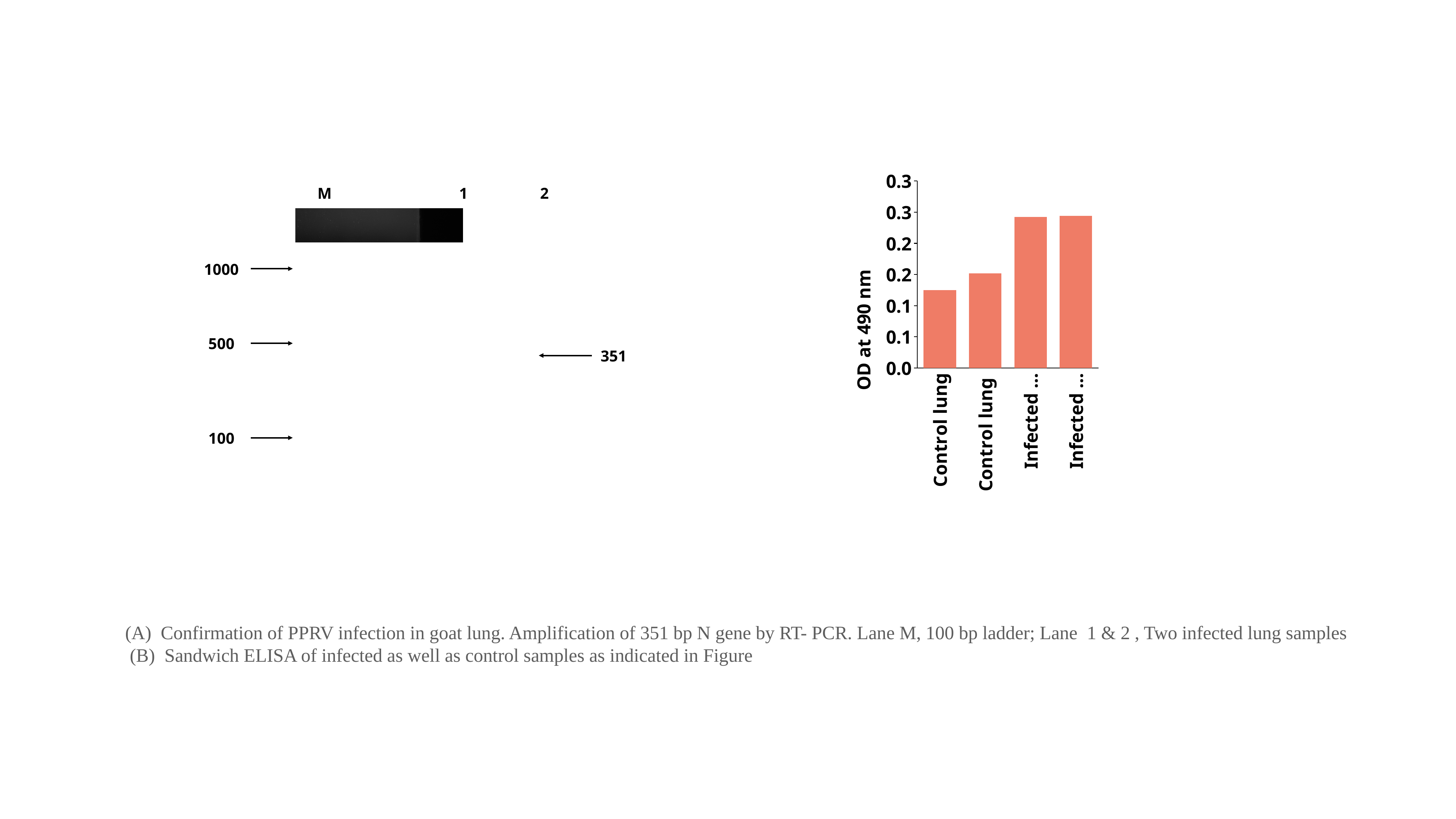

#### Chart
| Category | Untitled 2 |
|---|---|
| Control lung | 0.13 |
| Control lung | 0.158 |
| Infected lung | 0.252 |
| Infected lung | 0.254 |M
1
2
1000
500
351
100
(A) Confirmation of PPRV infection in goat lung. Amplification of 351 bp N gene by RT- PCR. Lane M, 100 bp ladder; Lane 1 & 2 , Two infected lung samples
 (B) Sandwich ELISA of infected as well as control samples as indicated in Figure
