## Supplementary Fille 5 for "Changes in m^6^A RNA methylation of goat lung following PPRV infection"

### Slide 1
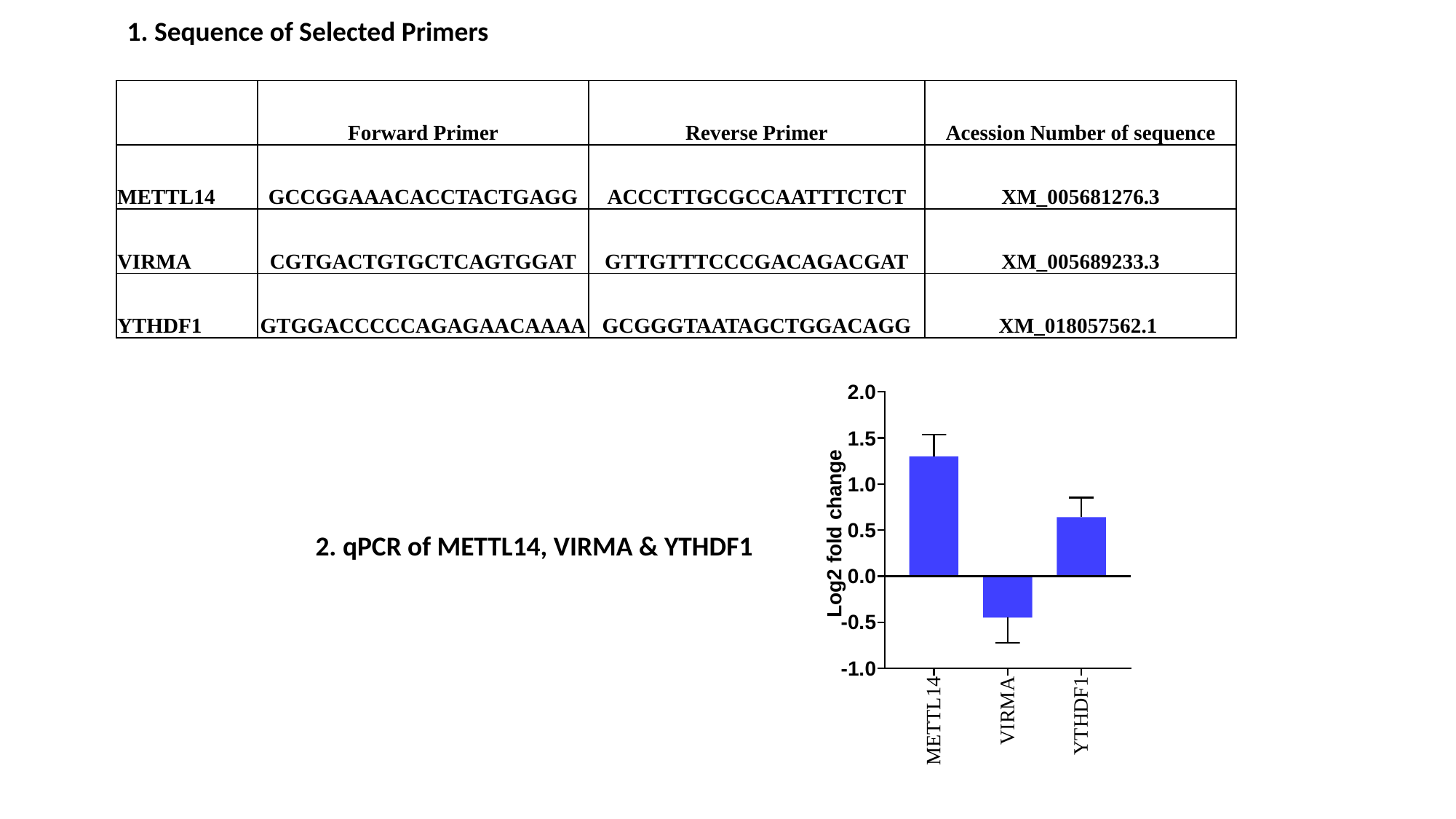

1. Sequence of Selected Primers
| | Forward Primer | Reverse Primer | Acession Number of sequence |
| --- | --- | --- | --- |
| METTL14 | GCCGGAAACACCTACTGAGG | ACCCTTGCGCCAATTTCTCT | XM\_005681276.3 |
| VIRMA | CGTGACTGTGCTCAGTGGAT | GTTGTTTCCCGACAGACGAT | XM\_005689233.3 |
| YTHDF1 | GTGGACCCCCAGAGAACAAAA | GCGGGTAATAGCTGGACAGG | XM\_018057562.1 |
2. qPCR of METTL14, VIRMA & YTHDF1
